# Commensal-pathogen dynamics structure disease outcomes during *Clostridioides difficile* colonization

**DOI:** 10.1101/2024.07.11.603094

**Authors:** Skye R. S. Fishbein, Anna L. DeVeaux, Sakshi Khanna, Aura L. Ferreiro, James Liao, Wesley Agee, Jie Ning, Bejan Mahmud, Miranda J. Wallace, Tiffany Hink, Kimberly A. Reske, Janaki Guruge, Sidh Leekha, Erik R. Dubberke, Gautam Dantas

**Affiliations:** The Edison Family Center for Genome Sciences and Systems Biology, Washington University School of Medicine, St. Louis, Missouri, USA; Department of Pathology and Immunology, Division of Laboratory and Genomic Medicine, Washington University School of Medicine, St. Louis, Missouri, USA; Division of Infectious Diseases; Washington University School of Medicine, St. Louis, Missouri, USA; Department of Molecular Microbiology; Washington University School of Medicine, St. Louis, Missouri, USA; Department of Biomedical Engineering; Washington University in St Louis, St. Louis, Missouri, USA; Department of Pediatrics; Washington University School of Medicine, St. Louis, Missouri, USA

## Abstract

Gastrointestinal colonization by *Clostridioides difficile* is common in healthcare settings and ranges in clinical presentation from asymptomatic carriage to lethal *C. difficile* infection (CDI). We used a systems biology approach to investigate why patients colonized with *C. difficile* have a range of outcomes. Microbiota-humanization of germ-free mice with fecal samples from toxigenic C. difficile carriers revealed a spectrum of virulence among clade 1 lineages and identified commensal *Blautia* associated with markers of non-pathogenic colonization. Using gnotobiotic mice engrafted with defined human microbiota, we observed strain-specific CDI severity across clade 1 strains. Yet, mice engrafted with a higher diversity community were protected from severe disease across all strains without suppression of *C. difficile* colonization. These results indicate that when colonization resistance has been breached without overt infection, commensals can attenuate a diversity of virulent strains without inhibiting pathogen colonization, providing insight into determinants of stable *C. difficile* carriage.

## Introduction

Healthcare-associated diarrhea caused by *Clostridioides difficile* remains difficult to predict, prevent, and treat, resulting in a significant and global public health burden^1^. While *C. difficile* infection (CDI) can be fatal, especially in the case of recurrence, *C. difficile* colonization in the absence of CDI (hereafter, *carriage*) is common. Recent studies in U.S. hospitals, by our group and others, have shown that toxigenic *C. difficile* carriage is more prevalent than symptomatic CDI, affecting approximately 10% of admissions in acute care and cancer-related cohorts^2,3^. Concerningly, these asymptomatic carriers are at substantially increased risk of developing future CDI, despite the possibility that these patients can also remain carriers without CDI for months at a time^2,3^. Despite the initial success of live biotherapeutics for recurrent CDI^4^, there is an unmet need to prophylactically intervene with such vulnerable patient groups, especially in a healthcare setting. Understanding what biological factors influence where patients lie along the spectrum from asymptomatic *C. difficile* carriage to CDI and identifying intervention targets is thus a public health priority.

The epidemiology of CDI has changed over the past two decades, with hypervirulent *C. difficile* lineages from clade 2 that were responsible for past epidemics (e.g., PCR ribotype 027/ST1^5^) no longer causing most healthcare-associated CDI. Genomic surveillance studies have instead found that clade 1 *C. difficile* strains (e.g., ST2, ST8, ST3, ST42)^2,3,6^ are predominant among currently circulating lineages isolated from both patients with CDI and asymptomatic carriers in multiple regionally distinct healthcare settings in the U.S.^2,3,6,7^. *C. difficile* is genetically diverse^16^, with strains showing marked differences in toxin production and metabolic preferences^8^. Quantifying the relative virulence of clinically prevalent *C. difficile* lineages, and their modulation by commensal microbes, is required to identify strategies aimed at preventing the transition from carriage to CDI in patients.

CDI is a paradigmatic disease of the gut microbiome, as loss of colonization resistance through microbiome perturbation often precedes CDI development^9^. Indeed, the remarkable efficacy of fecal microbiota transfers in preventing recurrent CDI depends upon restoration of colonization resistance^10^. Microbiota-conferred colonization resistance prevents *C. difficile* germination and vegetative growth through commensal bile acid, carbohydrate, amino acid, and micronutrient metabolism^11–14^. Contrastingly, certain commensal species such as *Bacteroides thetaiotaomicron* or *Enterococcus faecalis* have been shown to cross-feed *C. difficile*, supplying amino acids and carbohydrates to enable pathogenic growth and in some instances fuel disease onset^15–18^. The high prevalence of *C. difficile* carriage, where colonization resistance has been lost yet destructive toxigenesis is blunted, suggests that understanding these complex commensal-pathogen interactions may be key to preventing and treating CDI in vulnerable patients.

Validation of predicted commensal-pathogen interactions necessitates *in vivo* systems that successfully model patient microbiome structures while controlling for pathogen and commensal variation. Innovative gnotobiotic animal models using diverse defined communities have allowed us to understand the strain-specific nature of carbohydrate, amino acid, and bile acid metabolism in the gut microbiome, and such experimental systems would be ideal for gaining a deeper quantitative understanding of colonization resistance or resilience against *C. difficile*^19,20^. Further, it remains unclear how virulent circulating lineages of *C. difficile* are relative to earlier epidemic variants, complicating our understanding of how to pre-empt disease diagnostically or therapeutically. The identification of microbiome communities with desired resilience against diverse strains *in vivo*, and further modeling community dynamics using defined consortia, holds promise for examining how specific members of the microbiome can modulate the pathogenicity of *C. difficile*.

In this study, we use a systems biology approach to investigate how commensal-pathogen interactions influence the outcome of *C. difficile* colonization. We demonstrate how human gut microbiome structures impact *C. difficile* virulence by associating microbiome features with CDI status in colonized patients and identifying stable commensal-pathogen dynamics from patient microbiomes in microbiota-humanized gnotobiotic animal models. We present evidence for antagonistic roles of *Blautia* spp. in modulating *C. difficile* growth *in vivo* across a diverse collection of clinically relevant pathogen lineages. Importantly, our construction of defined gut microbial communities from patient stools allowed us to control for microbiome variation and identify that community structure can prevent toxin-mediated disease across these strains without inhibiting its colonization. These observations identify a dual role for the microbiome in supporting *C. difficile* growth while suppressing pathogenesis, which may have implications for the prevention of CDI in colonized patients and the design of novel microbiome-targeting therapies.

### Clinical and microbial features are discriminatory of CDI status in colonized patients

Identifying commensal, pathogen, and clinical features that render *C. difficile*-colonized patients vulnerable to CDI is essential to improving CDI treatment and prevention. Our previous examination of *C. difficile* carriers and patients with CDI revealed interactions between the microbiome, antibiotic exposure, and pathogen correlates that could drive patient outcomes^21^. We sought to investigate how these multilayered interactions underlie CDI risk in a broader group of hospitalized patients and controls. We leveraged stool samples and microbiome data from adult hospitalized cohorts between 2014 and 2019 of patients with CDI (CDI+, *C. difficile*+; *n* = 53) and *C. difficile* carriers (CDI-, *C. difficile*+; *n* = 96), with corresponding high-quality toxigenic *C. difficile* genome assemblies from 141/149 samples. We define CDI-as patients whose stool was either toxin enzyme immunoassay negative (EIA-) or not tested (based on a lack of clinical suspicion of CDI); toxigenic *C. difficile* carriage (*C. difficile*+) was confirmed by culture and/or stool nucleic acid amplification test (NAAT, see Methods for more detail). To contextualize the hospitalized patient microbiome, we additionally selected patient samples from these cohorts who did not have CDI symptoms and were *C. difficile* culture negative [hospitalized controls (HC); CDI-, *C. difficile*-; *n* = 39]. When available, we included longitudinal samples in our analysis and controlled for patient ID. Lastly, we extended our investigation to a control set of healthy volunteers without recent antibiotic exposure within the same geographic area (St. Louis, MO, USA)^22^ [healthy volunteers (HV); CDI-, *C. difficile*-; *n* = 20] (Fig. 1A). In total, we analyzed 208 stool metagenomes from 187 adults representing a spectrum of *C. difficile* colonization and disease status (Fig. 1A).

**Figure 1.**
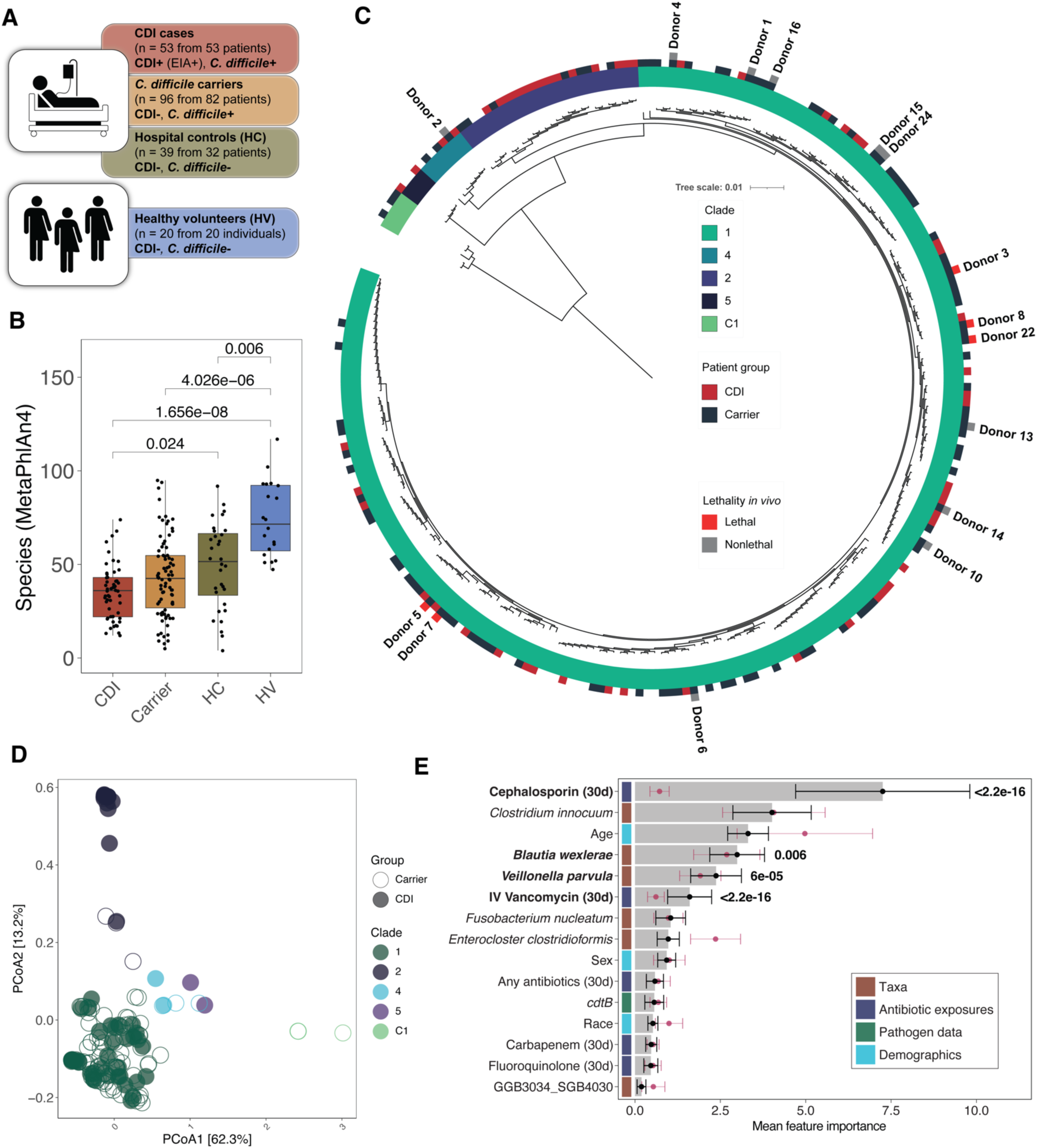
Microbiological variations underlie *Clostridioides difficile* colonization outcomes in a tertiary care hospital. A) Sampling scheme for human microbiome analysis including multiple hospitalized cohorts and a group of healthy volunteers within the same geographic area. B) Taxonomic richness of stool metagenomes (N = 53 CDI, 82 carrier, 32 HC, 20 HV) provided at the species level using MetaPhlAn4. Statistical tests performed using the Wilcoxon rank-sum test with Bonferroni correction for multiple comparisons. C) Phylogenetic tree (mid-point rooted) built from core gene alignment of our complete collection of *C. difficile* clinical isolates collected at Barnes-Jewish hospital. Inner ring corresponds to *C. difficile* clade (C1 = cryptic clade 1). Outer ring denotes isolates selected for this study and whether they derived from *C. difficile* carriers or patients with CDI. Isolates are labeled by donor ID for samples selected as microbiome engraftment donors in the following section and their lethality phenotype *in vivo*. D) Principal coordinate analysis of pan-genome alignment of selected *C. difficile* isolates plotted by Jaccard distance based on their pairwise average nucleotide identity (ANI). Each point corresponds to an individual genome and is colored by clade and filled based on CDI status. Percent variance explained is displayed parenthetically for the two axes shown. (C-D) depict data from 248 bacterial isolates. E) Mean feature importance plot based on a random forest classifier built to predict CDI status among colonized (carrier or CDI) samples using all features shown. The black and maroon points and standard deviation represent predictive importance of the empirical and null model, respectively. Features in bold are significantly more predictive than the corresponding null distribution (*p* < 0.05, Wilcoxon-rank sum test with Bonferroni correction for multiple comparisons). Color strip denotes feature type as referenced in the legend.

HVs had significantly increased taxonomic richness relative to carriers and patients with CDI, whereas HCs represented an intermediate group (Fig. 1B). This trend was also reflected by differences in species Shannon diversity (Fig. S1A) and microbiome structure as measured by beta diversity for HVs compared to all hospitalized patients (*p* = 1e-04, PERMANOVA; Fig. S1B). As many of these differences may be mediated by antibiotic exposure, we compared quantitative 30-day antibiotic exposures and identified significant differences in cephalosporin, fluoroquinolone, and intravenous (IV) vancomycin exposure across hospitalized patient groups (Table S1). Using ShortBRED^23^ to profile antibiotic-resistance genes (ARGs) in the microbiome, we observed an increase in both ARG burden and richness concomitant with *C. difficile* colonization and CDI, highlighting the substantial influence of antimicrobial exposure on differences in microbiome structure within this population (Fig. S1C-D). Finally, we integrated pathogen genomic information from the corresponding *C. difficile* strains of colonized patient samples^2^ and noted diverse representation of clades 1, 2, 4, 5, and cryptic clade 1 among our sample set, with a predominance of clade 1 among both carriers and patients with CDI (33/53 CDI cases and 68/78 carriers with cognate isolate assemblies; Fig. 1C)^6^, which matches other reports in the United States indicating the abundance of clade 1 in health-care settings^3,7^. Additionally, we observed that pangenomic differences in gene content among these selected strains clustered by clade (Fig. 1D). As previously observed^21^, patients colonized with strains encoding the accessory binary toxin, *cdtB*, were more likely to have CDI (*p* = 0.0028, Fisher’s exact test).

Because of these significant microbiome, antibiotic exposure, and pathogen correlates of CDI, we computed the relative importance of these variables for distinguishing between patients with CDI and asymptomatic carriers. *C. difficile* relative abundance is highly correlated with CDI status using linear mixed effect models (coef = 2.26, *qval* = 1.4e-07; Fig. S1E) and may overshadow the importance of other features; thus, we excluded it from downstream modeling tasks. We trained a random forest classifier on the relative abundance of feature-selected commensal taxa (*n* = 6 species), recent antibiotic exposures, participant demographic data, and the presence of *cdtB* for a randomized subset of colonized patients (60%) and assessed its accuracy in predicting CDI in the remaining data (40%) (see Methods). The highest performing model (of 100 iterations) had an accuracy of 84.3% with a positive predictive value of 80% and negative predictive value of 87.1% (Fig. S1F), demonstrating its effectiveness in discriminating between carriers and patients with CDI. Interestingly, in the absence of *C. difficile* relative abundance data, we observed that cephalosporins, IV vancomycin, *Blautia wexlerae*, and *Veillonella parvula* were all significantly important features in predicting disease status, while surprisingly, the mean feature importance of *cdtB* did not outperform those of models trained with randomly shuffled class labels (Fig. 1E). Exposure to cephalosporins or IV vancomycin and the abundance of *V. parvula* were all enriched in patients with CDI relative to carriers (Fig. S1E and Table 1), which corroborates the known influence of these antibiotic classes on CDI risk and identifying *V. parvula* as a commensal signature of CDI (Fig. S1E). In contrast, *B. wexlerae* was depleted in patients with CDI and negatively associated with recent cephalosporin exposure in colonized patients (Fig. S1E, G). In all, we identified specific antibiotic and commensal variables that were discriminatory of CDI status and found that *cdtB* was not an important feature despite its association with CDI. However, these insights also underscore the difficulty in identifying meaningful pathogen-microbiome interactions from metagenomic data, which are limited in ascribing functional importance to associations. Accordingly, we designed an *in vivo* microbiome engraftment screen to better understand the ecological determinants of *C. difficile* colonization dynamics.

### Microbiome engraftment screen reveals strain-specific drivers of commensal-pathogen dynamics

We hypothesized that we could leverage the natural microbiome-pathogen variation in patient stool samples to identify functional drivers of *C. difficile* proliferation *in vivo*. We devised a screen based on the premise that germ-free mice administered toxigenic strains of *C. difficile* succumb to lethal infection within 48 hours of gavage even at relatively low titers^12^. Thus, any non-lethal engraftment would reflect a stable equilibrium between the commensal microbiota and *C. difficile* (Fig. 2A). We selected 15 stools, or donors, from *C. difficile* carriers without CDI that reflect the ecological variation of colonized patient microbiomes and one stool from a HV (Donor HV), which lacks *C. difficile*, to engraft into germ-free mice (*n* = 3-8 mice per arm). We followed mice for 21 days post-engraftment to track disease, measuring clinical health, *C. difficile* burden, toxin production, and microbiome composition. Upon engraftment, we observed a spectrum of lethality (Fig. 2B), with 33% (5/15) of donor communities resulting in some lethality within the first week, while the rest (10/15) resulted in non-lethal engraftment. There were also community-specific differences in weight loss trajectories during the first week, with lethal donors exhibiting significant weight loss relative to Donor HV, which did not contain *C. difficile* (Fig. S2A). We first investigated how well these stool engraftment experiments modeled human microbiome ecologies in the context of *C. difficile* colonization. Comparison of human microbiome structures of the selected stools revealed that the *C. difficile* burden of the original donor stools correlated with distance (as measured by beta diversity) from the HV cluster, where no *C. difficile* was detected (Fig. 2C, D; Pearson *R* = 0.76, *p* = 0.0011), suggesting that a higher degree of dysbiosis correlates with *C. difficile* abundance. While there was significant correlation (Spearman *R* = -0.71, *p* = 0.0031) between donor lethality in the first week and *C. difficile* colony-forming units (CFU)/mL in the inoculum gavage (Fig. S2B), one donor (Donor 3) caused lethal disease despite a relatively low *C. difficile* inoculum titer. At 21 days, mice from donors surviving initial engraftment (thus excluding donor 5 and 7) had a community species richness that fell within the range of colonized human microbiome richness (species richness of colonized patient microbiomes ranged from 12-74; Fig. 2E). To investigate *C. difficile* fitness and virulence in non-lethal donor groups, we compared *C. difficile* CFUs and TcdB levels in mouse stools at an early point (days 4-5) and a late timepoint (day 21) and calculated ratios of *C. difficile* growth and toxin production throughout engraftment (where a value above 1 indicates an increase in pathogen abundance or toxin levels over time), revealing donor-specific trajectories in *C. difficile* proliferation and toxin levels post-engraftment (Fig. S2C, D).

Our human microbiome-genomic data suggested that microbiome structure and not *C. difficile* strain identity drives symptomatic severity. In this functional microbiome screen, we sought to reexamine the contribution of these two factors to longitudinal differences in *C. difficile* growth and virulence following stool microbiome engraftment. Further, given our and others’ observations that clade 1 isolates are the major circulating strains in the United States^2,3,6,7^, our microbiome engraftment screen contained mostly stools with *C. difficile* strains from this clade (14/15 stools), including both closely related isolates and unrelated isolates clusters, based on whole-genome information (Fig. 2F). To understand strain-specific influences on outcomes in our microbiome engraftment screen (weight-loss, *C. difficile* titers, and toxin levels), we aggregated clade 1 donor strains into groups described by their multi-locus sequence type (sequenceT), based on a threshold of >99.9% ANI by core-gene alignment of selected clade 1 genomes. Examination of weight-loss differences revealed that stools containing ST8 and ST43 strains caused significantly more weight-loss relative to other strains and the ST2 strain group (Fig. 3A). Though some stools containing these two strain groups had high levels of *C. difficile* in the gavage inoculum, Donor 3 contained a low level of its ST8 strain (Fig. S2B), suggesting that this strain group may be more virulent than other clade 1 isolates examined here. To examine strain influence on growth and toxin production, we restricted our measurements to strain groups with multiple non-lethal stool microbiome backgrounds. Interestingly, the ST2 strain group had decreased *C. difficile* proliferation and toxin production over time relative to *C. difficile* strains in all other donors, suggestive of lower virulence independent of the microbiome (Fig. 3B, 3C). Juxtaposed with the increased lethality associated with ST8 strains from Donor 3 and Donor 8, our microbiome engraftment screen revealed significant variation in clade 1 strain fitness and toxin production *in vivo*.

**Figure 2.**
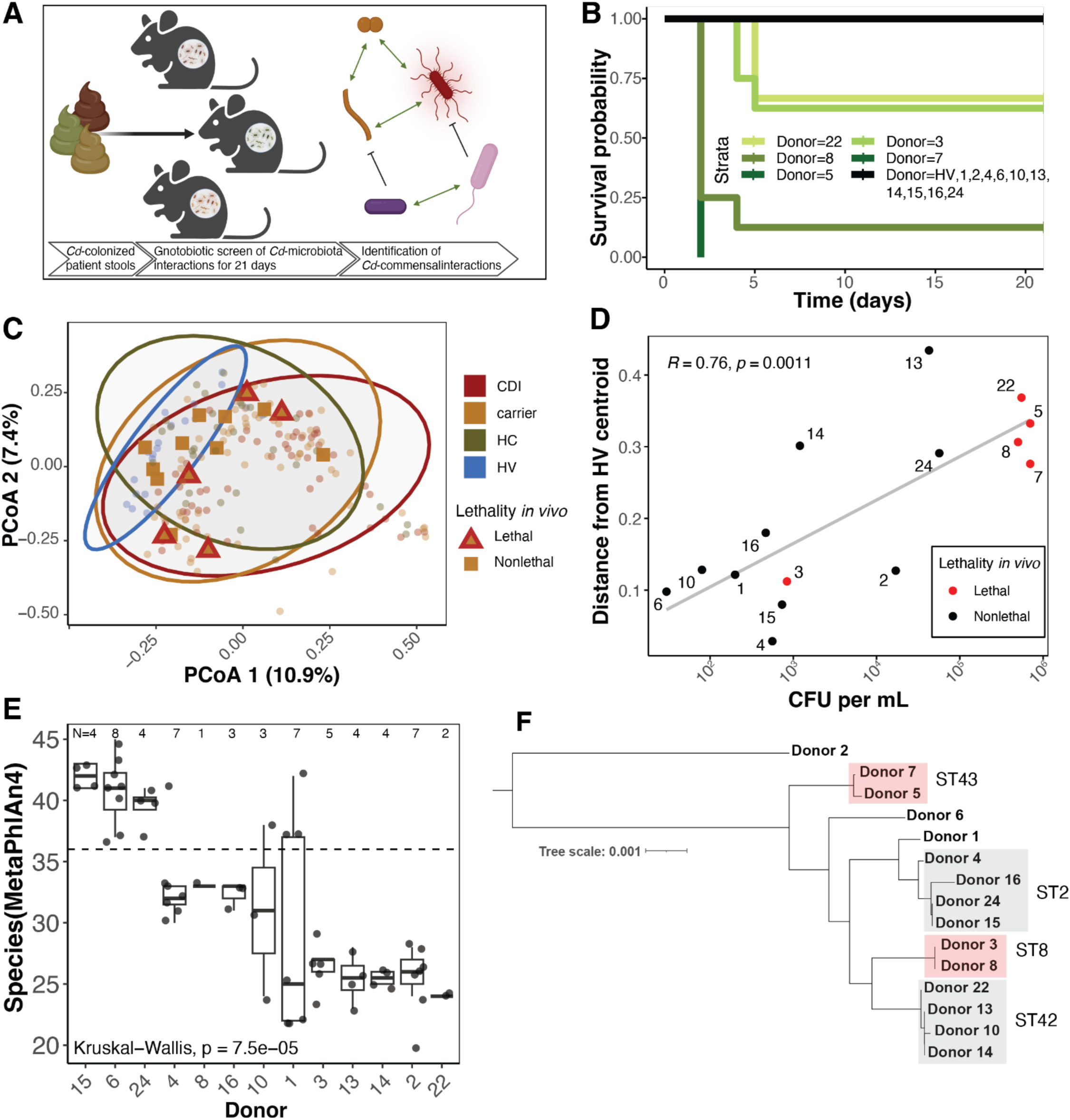
Microbiome engraftment screen recapitulates the spectrum of human *C. difficile*-associated disease. A) Humanization of germ-free mice with carrier patient microbiomes allows observation of pathogen-microbiome dynamics. B) Kaplan-Meyer plot of microbiome humanization, where each color reflects a stool (Donor) that caused >0% lethality in mice (*n* = 3-8 mice per group, see Methods). C) Principal coordinate analysis of Bray-Curtis dissimilarity across human microbiomes (●), where donors used in the screen are indicated as lethal (▴) if any mice died or nonlethal (▪). (▪) (*n* = 53 CDI, 83 carrier, 32 HC, 20 HV). D) Relationship between microbiome community structure and donor stool *C. difficile* abundance (CFU/mL), as measured by distance from HV communities in C. *R-*squared value and p-value from a Pearson correlation test. E) Richness of microbiomes in surviving mice at day 21 (N indicates number of mouse stool or fecal microbiome quantified for each donor), as measured by MetaPhlAn4 species. F) Midpoint-rooted phylogenetic tree built from core gene alignment of 15 *C. difficile* isolates associated with donor stools. Boxes indicate a relatedness of >99.9% ANI; grey boxes indicate isolates associated with non-lethal donors while red boxes indicate >0% lethality associated with donor stools.

**Figure 3.**
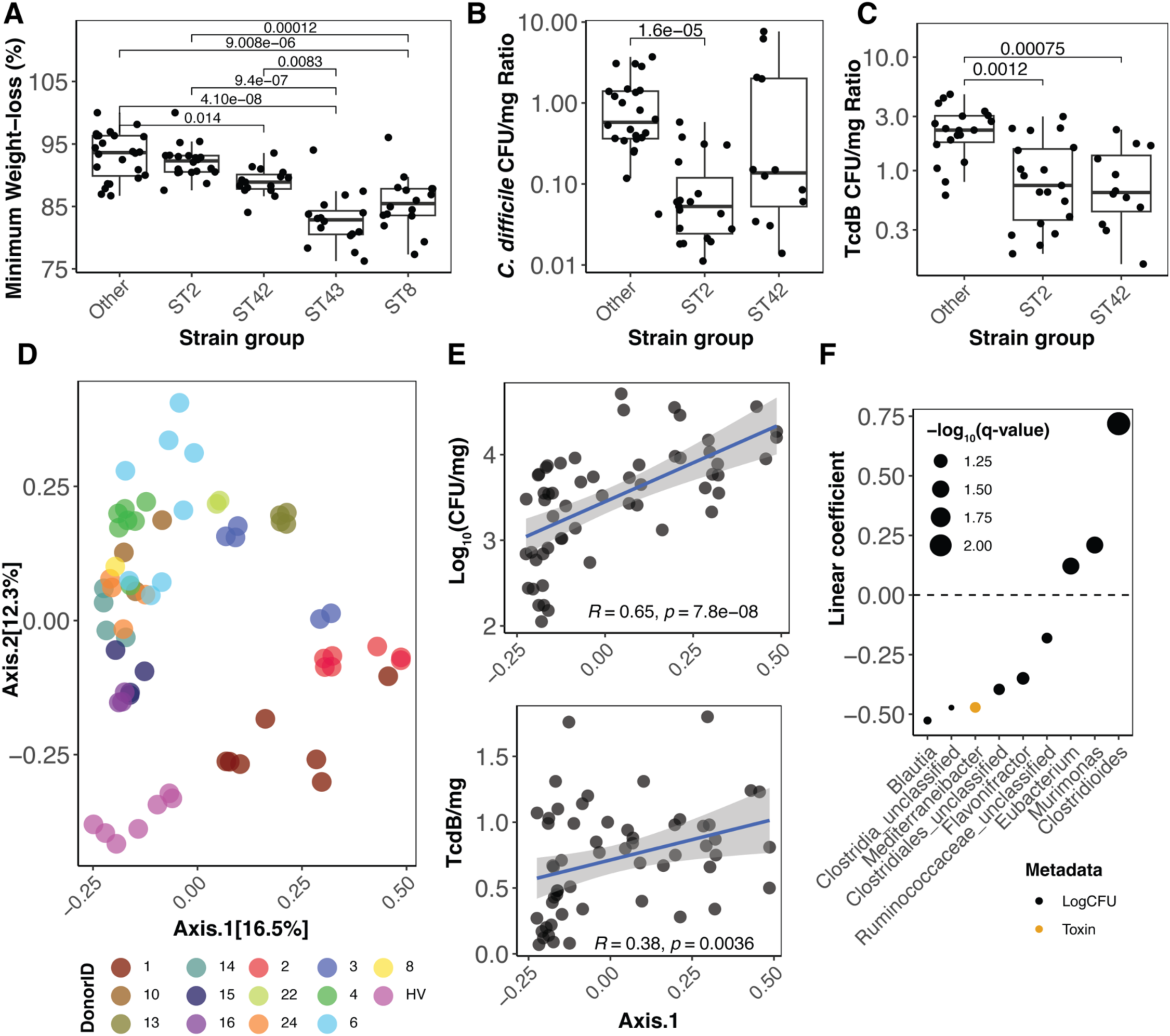
Strain variation and microbiome structure drive differences in colonization outcome *in vivo*. *C. difficile* strain groups display significant differences in A) weight-loss, B) titer ratios, and C) toxin/mg ratios. P-values indicate Benjamini-Hochberg corrected results of a Dunn’s test comparison between mouse measurements across different strain groups, and *n* indicates the number of mice from which ratios was calculated. D) Principal coordinate analyses of Bray-Curtis dissimilarity between all fecal microbiomes of surviving mice; each point represents a stool microbiome from one mouse timepoint at day 21. E) Relationship between microbiome structure (PCoA from [D]) and stool *C. difficile* (top) and toxin levels (bottom). *R* value and p-value indicate the results of a Pearson correlation test. F) Linear coefficients of taxa related to growth or toxin levels in mice with a q-value <0.1 using MaAsLin2 as described in the Methods (analyzed 88 stool metagenomes over time across 13 donor backgrounds).

We leveraged the above pathogen genomic information to more precisely define the commensal contribution to *C. difficile* proliferation and toxin production in our screen. Using beta diversity across all sequenced murine microbiomes to generate a principal coordinate analysis, we confirmed human donor-specific clustering of gnotobiotic mouse microbiome structures (Fig. 3D) and identified a significant correlation between gnotobiotic mouse microbiome structure at day 21 and corresponding mouse stool levels of *C. difficile* (Fig. 3E, *R* = 0.65, *p* < 0.001) and, to a lesser extent, its toxin (Fig. 3E, *R* = 0.38, *p* = 0.0036). We then used linear mixed effect modeling to investigate taxonomic features that were associated with quantitative differences in *C. difficile* growth in stool while controlling for *C. difficile* strain identity and other microbiome-relevant factors (see Methods). Importantly, our screen revealed that *Blautia* was strongly associated with decreased *C. difficile* growth (Fig. 3F), validating correlative data from our human microbiome analyses indicating that *Blautia wexlerae* depletion (perhaps related to antibiotic exposure) was a signature of CDI in patient microbiomes (Fig. 1E, S1E). Based on these metagenomic findings from both patient samples and gnotobiotic colonized communities *in vivo*, we predict that *Blautia* spp. interact with *C. difficile* during stable colonization and modulate its growth. We further find that several carbohydrate catabolism pathways, including glycogen degradation and three proteoglycan-related sugar acid degradation pathways, were associated with increased *C. difficile* growth (Fig. S2E). These taxonomic and metabolic pathway associations indicate that *Blautia* and complex glycan metabolism may be involved in stabilizing *C. difficile* during non-pathogenic colonization *in vivo*.

### Defined microbiome enables *C. difficile* colonization and suppresses strain-specific virulence

While we identified multiple correlates of *C. difficile* proliferation during colonization *in vivo*, including both strain identity and microbiome composition, we sought to further understand strain differences in the globally prevalent clade 1 and identify strategies by which the microbiome alters *C. difficile* virulence in a controlled manner. To achieve this, we built two defined bacterial culture collections representative of patients carrying *C. difficile* in our cohort as a background for subsequent *C. difficile* challenge *in vivo*. These defined communities were designed to either 1) recapitulate the phenotype of a candidate non-lethal donor in our microbiome engraftment screen or 2) broadly represent the most prevalent taxa seen in *C. difficile* colonized patients based on metagenomic analysis. Thus, for the first defined community (DefCom1), we sought to reassemble the community (except for *C. difficile*) from donor 4, for which we observed intermediate weight loss *in vivo* (Fig. S2A); DefCom1 comprised 24 species representing 98% of the relative abundance of species observed in the original donor stool (Supplementary Table 2). For the second community (DefCom2), we supplemented DefCom1 with additional taxa to represent at least 70% of the 30 most prevalent genera among colonized patients, totaling 46 commensal species (Fig. 4A). We pooled each set of isolates and engrafted germ-free mice with two consecutive daily gavages of either the DefCom1 or DefCom2 defined culture collection. After 28 days, both communities were representative of human microbiome states found in colonized patients (Fig. S3A).

**Figure 4.**
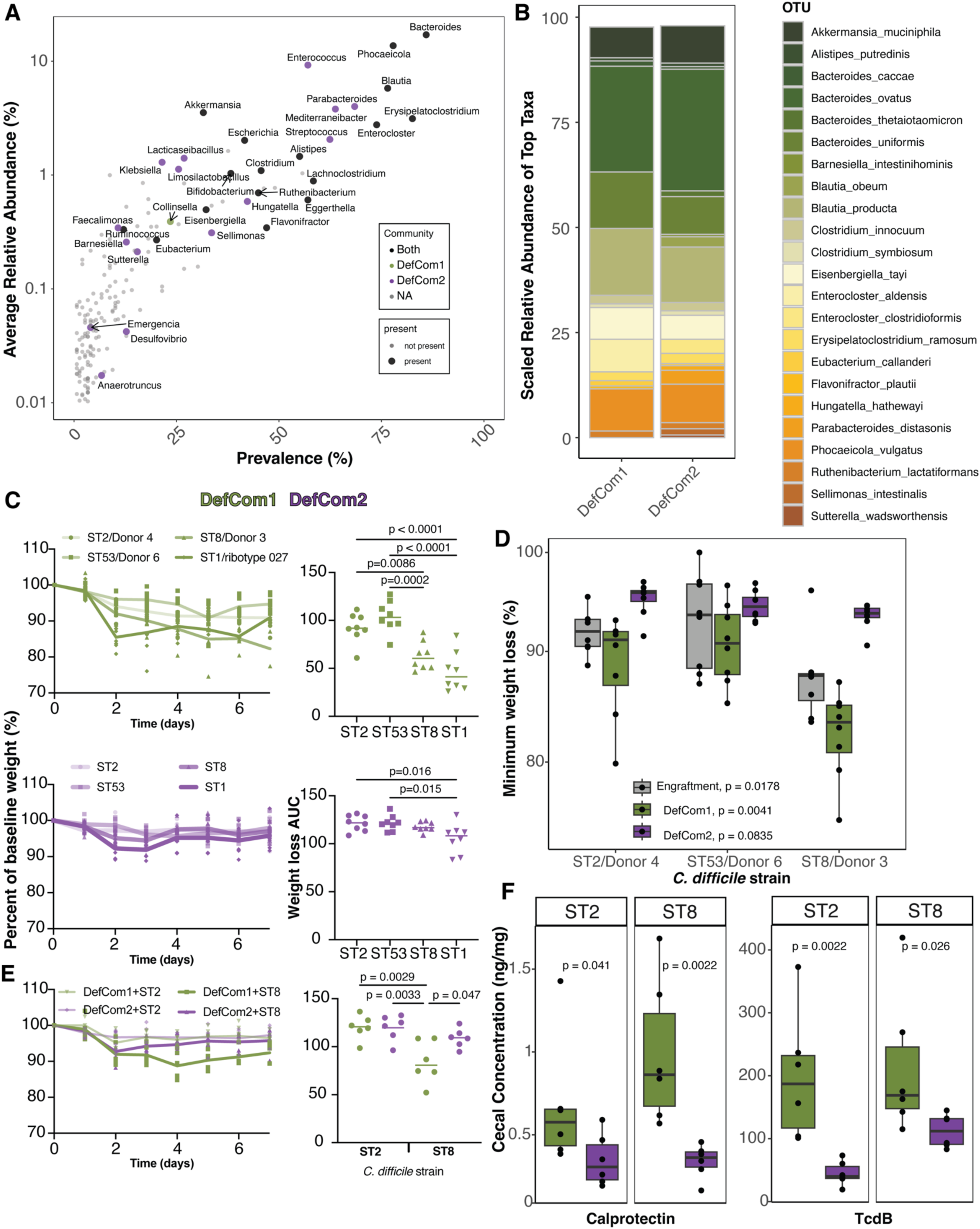
Construction of defined human gut microbiota in gnotobiotic models recapitulates spectrum of *C. difficile*-associated disease. A) Prevalence and abundance of genera in patients colonized with *C. difficile* from Figure 1, where color indicates use in DefCom1 or DefCom2. B) Metagenomic abundance of species at >0.1% relative abundance in DefCom1 and DefCom2 (representative of average abundance across DefCom1 [N=43 mouse stool metagenomes] and DefCom2 [N=52 mouse stool metagenomes] at day 28 in all experiments). C) Weight-loss after infection with *C. difficile* isolates following engraftment in DefCom1 (top, N=8 mice in each arm) and DefCom2 (bottom, N=8 mice in each arm). D) Minimum weight-loss in the first week after engraftment (from Figure 2, N= 8 mice per donor arm) or infection from (C). P-values indicate Kruskal-Wallis test comparison of weight-loss within a single microbiome background. Weight-loss percentage following infection with least virulent (ST2) and most virulent (ST8) clade 1 isolates, repeat of (C); N=6 mice per arm. F) Cecal levels of *C. difficile* toxin and calprotectin levels across *C. difficile* strains and microbiome backgrounds at day 7 post-infection from (E). P-values indicate Wilcoxon rank-sum test performed across different microbiome backgrounds for the same *C. difficile* strain. For (C) and (E), one-way ANOVAs were performed with Tukey’s multiple comparisons test with a single pooled adjustment.

To investigate differences in virulence among the diverse clade 1 lineage, we chose three clade 1 strains from our engraftment experiments that reflected differences in mouse weight loss, lethality, and *C. difficile* growth. Specifically, we selected strains natively colonizing donor 4 (the source community for DefCom1) from the least ‘virulent’ strain group (ST2, Fig. 2D, F); donor 6 (ST53), which had the highest TcdB load in non-lethal donors; and donor 3 (ST8), for which we observed intermediate lethality at a relatively low *C. difficile* load and thus predicted higher virulence (Fig. 2D). Following engraftment with DefCom1 or DefCom2, mice were challenged with 10^4^ CFU of ST2, ST53, ST8, or a ribotype 027 reference strain (ST1, UK1)^24^. Relative to the hypervirulent ST1 infection, we observed strain-specific virulence of clade 1 isolates (Fig. 4C) across both communities, with ST8 resulting in intermediate disease and ST2 and ST53 causing mild disease. Notably, DefCom1 colonization resulted in more severe disease, as measured by lethality and weight loss, upon *C. difficile* challenge for all strains relative to DefCom2. DefCom2 suppressed strain-specific differences in weight loss relative to microbiome structures in both DefCom1 and human fecal engraftment backgrounds (Fig. 4D, data from Fig. S2); interestingly, this effect was most pronounced for ST8, the most virulent clade 1 isolate.

We sought to validate the observed difference in microbiome protection between DefCom1 and DefCom2 using ST2 and ST8 strains in an independent experiment; a comparison of microbiome engraftment at Day 28 indicated that we reproduced community structure between experimental replicates (Fig. S3B, C). In addition to recapitulating weight loss severity differences (Fig. 4E), we found that DefCom2 resulted in decreased toxin production and decreased intestinal inflammation (as measured by calprotectin, which is induced during CDI ^25,26^) relative to DefCom1 for both strains (Fig. 4F). Surprisingly, for the moderately virulent ST8 strain, DefCom2 supported a higher level of *C. difficile* as measured by CFUs (Fig. S3D) relative to DefCom1. To rule out the possibility that absolute bacterial loads explained the difference in ST8 *C. difficile* CFU load between communities (i.e., carrying a similar abundance of *C. difficile* relative to community absolute abundance), we plated pre-challenge (day 28) stools from each repeat experiment on LYBHI with sodium taurocholate to recover the majority of spore and vegetative cells in the community. Interestingly, we observed equivalent bacterial load between DefCom1 and DefCom2 after two days of growth (Fig. S3E), suggesting preferential growth of ST8 in DefCom 2 background.

Given the results from our microbiome engraftment screens, we postulated that the community-specific difference in disease severity and the promotion of ST8’s growth by DefCom2 reflects the microbiome’s capacity to cross-feed *C. difficile*, thus encouraging its proliferation while attenuating virulence. Specifically, based on the well-studied repression of toxin production in the presence of metabolizable nutrients^27^, we predict that the increased flux of host- and dietary-derived nutrient degradation products released by commensals represses *C. difficile* toxin production. Investigation of metagenomically-inferred metabolic pathway differences between DefCom1 and DefCom2 predominantly revealed an enrichment in glycan degradation pathways, including that of host-derived glycosaminoglycans and plant polysaccharides (Fig. S3F). These associations support a potential role for glycan metabolism in promoting *C. difficile*-commensal interactions during stable colonization, which may contribute to its attenuated virulence across Clade 1 strains in DefCom2.

## Discussion

*C. difficile* 630 (a clade 1 isolate of ST54^28,29^) and a number of well-studied clade 2 strains^29–31^ are genetically tractable, and these strains have enabled a foundational understanding of how this bacterial pathogen grows and causes disease^29^. Importantly, previous work in a conventional antibiotic-induced CDI murine model has demonstrated differential virulence between *C. difficile* 630 and these epidemic strains, recapitulating both CDI and non-pathogenic colonization^32^. However, ribotype 012, which includes *C. difficile* 630, is an uncommon lineage identified in patients today and may not be sufficiently representative of other clade 1 strains in terms of studying *C. difficile* colonization outcomes. Given our growing understanding of the vast diversity of *C. difficile* lineages (owing to its open pangenome^6^), we sought to expand our understanding of clade 1 virulence in the context of gnotobiotic models of patient-derived microbiomes. Our data show that, across two microbiome backgrounds, only one of the clade 1 isolates tested (a strain from ST8) causes levels of initial weight loss in mice equivalent to those of the hypervirulent ST1 from clade 2 (Fig. 4). These data are consistent with previous reports that ribotype 002, to which our ST8 strain belongs, may have significant virulence relative to other common circulating clade 1 lineages^33,34^. In contrast, the ST2 strain produced minimal weight loss, supporting the trend of overt differences in virulence that others have observed between these clades of *C. difficile* and underlining the variation in clade 1 virulence^32^. Notably, ST8 and ST2 are the most globally prevalent clade 1 lineages of *C. difficile*^6^. Although ribotype 027 (clade 2) was still the single most common ribotype identified in patients with healthcare-associated CDI in 2018, from an epidemiologic perspective, it appears that historically less virulent strains are the predominant source of *C. difficile* carriage and CDI. Future investigation of how clade 1 relative to other clade 1 and clade 2 isolates cause severe disease in equivalent microbiome backgrounds is warranted to both adapt surveillance strategies in vulnerable patient populations and to improve our fundamental understanding of *C. difficile* pathogenesis across a greater representation of clinically and epidemiologically significant pathogen genomic diversity.

We identified an inverse correlation between *Blautia* spp. and CDI in colonized patients and with *C. difficile* growth in our microbiome engraftment screen. *Blautia* has previously been implicated in human and animal microbiome-association studies as significantly altered in *C. difficile*-related disease outcomes^35–37^, primarily associated with protection from disease. Consistent with such findings, these taxa appear to blunt *V. cholera* proliferation through bile acid metabolism ^38–41^. More recent genome-scale metabolic modeling indicates that *Blautia* may compete with *C. difficile* for amino acid pools^42^. These studies suggest that in our microbiome engraftment screen, *Blautia* spp. either alter germination rates of *C. difficile* or slow down vegetative growth by consuming overlapping nutrient pools. Additionally, data from our clinical cohort indicates several Lachnospiraceae to be depleted in patients with recent cephalosporin exposure, which was most indicative of CDI in colonized patients, underlining the clinical significance of these ecological relationships.

Gnotobiotic animal models of *C. difficile*-commensal interactions have made critical progress in illuminating the mechanisms by which commensals support *C. difficile* proliferation through carbon-source crossfeeding^12,18^ or inhibit *C. difficile* disease through competitive Stickland fermentation^12^ and secondary bile acid generation^25^. These approaches relied on simple communities (*n* = 1-13 commensal species) to profile the transcriptional and metabolomic landscape of *C. difficile* colonization *in vivo*^12,18,25^. More recently, a gnotobiotic model using three gut commensals revealed that *C. difficile* consumption of host nutrients decreased virulence^43^. Based on these approaches, we hypothesized that we could model diverse human microbiome variation that would help explain differences in CDI susceptibility among colonized patients and provide a controlled background for assessing the virulence of clinically prevalent *C. difficile*. The defined culture collection approach allowed us to recapitulate *C. difficile* carriage, where gnotobiotic animals colonized with DefCom2 attenuated the strain-specific differences in weight loss (among clade 1 and 2 isolates) seen with DefCom1. DefCom2’s distinct suppression of toxin production and toxin-mediated inflammation across diverse strains (from clade 1 and 2) indicates that, in a given patient, certain microbiome structures that enable *C. difficile* colonization may overcome strain-specific virulence attributes. Further prediction of pathway utilization in these different microbiomes reveals that DefCom2 is enriched in polysaccharide/sugar degradation and glycosaminoglycan degradation. Given that the ST8 strain grew to higher abundance in the DefCom2 background, we speculate that DefCom2 feeds *C. difficile* host- and/or diet-derived glycan degradation products and thus represses cellular stress and subsequent toxin production. This hypothesis fits with our longstanding understanding of nutrient-mediated transcriptional repression of toxin production^8,44,45^ and is supported by recent data indicating that commensal competition may increase toxin production^42^. These defined microbiome data in complex with quantitative measurements of strain virulence indicate that *C. difficile* is not exclusively an acutely destructive gastrointestinal pathogen but can instead operate as a pathobiont capable of cooperating in a complex microbial community.

We acknowledge the limitations of our study and highlight opportunities for future work to further examine how commensals can support stable *C. difficile* colonization and suppress virulence. Our human cohorts were derived from a single healthcare setting in the U.S., and while our observed toxigenic *C. difficile* colonization rates and the prevalence of clade 1 isolates^2^ mirror other recent reports in Michigan^3^ and Massachusetts^7^, it will be important to extend such analyses outside the U.S. and beyond the healthcare setting. It is notable that the patients who served as donors for engraftment experiment did not have clinical CDI, yet a fraction of those donor stools produced severe disease in mice, suggesting that not all species present in the donor engrafted into mice. While our defined communities were rationally designed to be highly representative of microbiome states observed in stably colonized patients, they are not inclusive of all taxa found in these communities, and the construction of additional consortia or augmentation of our existing communities may reveal differences in resilience against CDI, such as the taxa that are most affected by antibiotic exposure. Although we identify DefCom2 to attenuate disease from virulent clade 1 and clade 2 *C. difficile* strains compared to DefCom1, strain dropout experiments^19^ will be crucial for systematically assigning which taxa are necessary and sufficient for such resilience and in which combinations, and, lastly, whether this protection extends to other clinical *C. difficile* lineages (such as the globally prevalent ST11 from clade 5)^6^.

Our work highlights the provocative potential for future anti-CDI therapies which could suppress toxin production while still supporting asymptomatic *C. difficile* growth and colonization in the gut. If surveillance approaches continue to report stable colonization as a defining feature of circulating *C. difficile* lineages, it will be important to better predict disease based on microbiome structure and relevant antibiotic exposure. Furthermore, the development of novel microbiome therapeutics that specifically focus on suppressing toxin production are warranted, with concomitant consideration of strain-specific features of the commensal microbiome that cooperate with or antagonize pathogenic proliferation and epithelial destruction.

## Methods

### Integration of human microbiome samples

Stool samples were collected from inpatients at Barnes-Jewish Hospital (BJH) in St. Louis, MO, USA in two cohorts (retrospective and prospective)^2,46^. Stool specimens that were submitted for toxin enzyme immunoassay (EIA) testing were collected from patients in the retrospective cohort who had clinically significant diarrhea and no alternate identifiable cause of diarrhea from 2014-2016 as previously described^46^. Samples that were metagenomically sequenced to sufficient depth and had corresponding toxigenic *C. difficile* genome assemblies were included here for analysis for a total of 48 primary CDI episode (EIA+) and 54 carriage (EIA-) samples^21^. Microbiome and isolate assembly data were downloaded from BioProject accession numbers PRJNA748262 and PRJNA980715, respectively. The prospective cohort was drawn from a surveillance study of patients admitted to the BJH hematopoietic stem cell transplant and leukemia units^2^ during a four and six month period, respectively, in 2019. Stool and/or rectal swab samples were collected at the time of admission and weekly thereafter and were cultured as previously described^2^. As with the retrospective cohort, specimens submitted to the BJH microbiology laboratory for EIA testing due to clinical suspicion of CDI were cultured for *C. difficile*. In the prospective cohort, carriers were classified as being culture positive for toxigenic *C. difficile*, did not develop CDI within 30 days of stool collection date, had no documented CDI episodes in the previous 8 weeks, and were EIA-in the case of suspected CDI. One patient-timepoint was flanked by two positive toxigenic cultures and both *C. difficile* and the *tcdA/B* genes were detected in the stool metagenome, so the patient was also classified as a carrier. Hospitalized control samples were within 7 days of a *C. difficile* culture-negative sample and did not become culture-positive during hospitalization; furthermore, samples for which *C. difficile* was detected metagenomically were excluded. Samples of patients with CDI (EIA+) were selected if they were stool specimens submitted for EIA testing (i.e., before CDI treatment initiation) and represented a first episode of CDI (i.e., no documented CDI within the previous 8 weeks). Longitudinal samples, which applied to a subset of carriers and hospitalized controls from the prospective cohort, were included as available for linear mixed effect modeling (which controlled for patient ID, see below) and one sample per patient was randomly selected as the “primary” sample for all other analyses (for one patient, two samples were included in the PCoA visualizations as they were both used as donor stools). For this cohort, *C. difficile* genome assemblies were downloaded from BioProject accession number PRJNA980715; a core-genome alignment was constructed using panaroo and fastree was used to construct a phylogenetic tree, as previously^47^. Antibiotic exposures in the 30 days prior to stool collection were encoded as binary variables and compared between patient groups using a Fisher’s exact test for subclasses with at least 10% prevalence across both hospitalized cohorts. Healthy volunteer microbiome data was drawn from a previously published cohort of adult individuals in the same geographic area without recent prior antibiotic exposure^22^; sequencing data was downloaded from BioProject PRJNA664754.

### Metagenomic sequencing and analysis

Metagenomic DNA was extracted from patient stool from the prospective cohort and mouse fecal matter (for all animal experiments) using the Qiagen PowerSoil Pro kit. Libraries were prepared using a modified Nextera protocol as previously^21^ and sequenced as 2×150 paired-end reads on an Illumina NovaSeq 6000 or NextSeq 550 instrument. Reads were demultiplexed by their index pair and quality-trimmed using Trimmomatic^48^ (v0.39, SLIDINGWINDOW: 4:20, LEADING: 10, TRAILING: 10, MINLEN:60) followed by decontamination of human or mouse genomic reads using DeconSeq^49^(v4.3; -dbs hsref38 or mm_ref_s1 & mm_ref_s2, respectively). MetaPhlAn4 (mpa_vJan21_CHOCOPhlAnSGB_202103) database was used to profile taxonomic composition and lowly abundant taxa were removed by filtering out species calls below 0.1% relative abundance^50^. HUMAnN3 was used to profile community metabolic function and normalization of pathways was performed to relative abundance^50^. Antimicrobial resistance gene (ARG) profiling of human metagenomes was performed using shortBRED against a marker gene set built from the NCBI and CARD databases^23^. ARG burden was determined using reads mapping per kilobase per million. Rarefaction was performed to identify sequencing thresholds for human microbiomes or mouse microbiomes based on subsampling a random set of ten samples from nine million paired-end reads to 1000 reads using seqtk (https://github.com/lh3/seqtk). Subsampled metagenomic reads were profiled using MetaPhlAn4 to assess taxonomic diversity, and differences between species richness were assessed for significance using a Wilcoxon rank sum test. Correspondingly, samples for human analyses that accrued less than 2.5 million paired-end reads per metagenome were removed from further analyses. Samples for mouse analyses that accrued less that 2 million paired-end reads per metagenome were removed from further analyses. Pathways below 0.01% relative abundance or assigned as ‘UNINTEGRATED’ were removed. For comparison of DefCom1 and DefCom2 functional pathways, aggregation of pathways was performed using a MetaCyc “parent orthology” classification^51^.

### Random forest classifier

Random forest classifier analyses were carried out as described previously^52^ in R with modifications using caret v6.0.94. The dataset consisted of all colonized patient samples with corresponding high-quality *C. difficile* genome assemblies and available race information for the participant; the “primary” sample was selected for a given patient (n = 128 [50 CDI, 78 carriers]). This dataset was randomly split into training (60%) or testing (40%) subsets (seed = 42). Taxonomic features were selected with Boruta (v8.0.0) using species relative abundances from the training data (after excluding *C. difficile*) and identifying taxa selected in at least 25 of 100 iterations (seeds 1:100). The six selected taxa were subjected to preprocessing using the caret “center” and “scale” transformations, while 30-day antibiotic exposures (>10% prevalence), demographic data, and *C. difficile* binary toxin (*cdtB*) presence or absence were encoded as categorical variables. Random forest classifiers were trained on 100 random subsets of 80% of the training cohort using a different seed in each instance (seeds 1:100) with 10-fold cross-validation in each case using the “Accuracy” metric. Each model was tested against the retained validation cohort and the most accurate model was used to summarize model performance. To determine feature importance, null models were trained on datasets with randomly shuffled class labels and the resulting feature importances were determined using the caret varImp function with scale = FALSE. The mean feature importance per variable was determined for each model tested and all empirical and null distributions were compared using Wilcoxon rank sum test and Bonferroni correction for multiple hypotheses.

### Microbiome engraftment screen

Stools were selected from carriers in this study and used for fecal microbiota engraftment of germ-free mice under Washington University in St. Louis IRB #202203020. Gnotobiotic mouse experiments were performed using protocols approved by the Washington University School of Medicine Institutional Animal Care and Use Committee (21-0160). Adult male and female germ-free mice were housed in Allentown SPP Sentry racks and fed Lab Diet 5K67 autoclavable chow with a 12 hour on, 12 hour off 6am – 6pm light cycle. Stools from donors HV, 1, 2, 3, 4, 5, 6, 7, 8, and HV were engrafted into equal numbers of male and female mice (N=8) over the course of two experiments. Stools from donors 10, 13, 14, 15, 16, 22, and 24 were engrafted into equal numbers of male and female mice (N=4) over the course of one experiment; one mouse in the donor 22 arm suffered a lethal cecal torsion before the experiment was initiated, thus decreasing the number of mice (N=3). To prepare stools for gavage or culturing, fecal matter was resuspended in PBS-30% glycerol-0.5% cysteine and filtered through a 100uM filter (Falcon) to remove debris. Filtered gavage material was stored at -80 degrees C until gavage. Mice were gavaged with 100uL of gavage material on two consecutive days. Following gavage, mice were monitored for weight-loss and clinical signs of disease (marked by hunched posture and slowed movement)^53,54^. Upon 15% weight-loss (relative to weight on the first day of gavage) and or severe clinical disease (moribund), mice were euthanized. Stool was collected at least once per week following engraftment, and each week following through 21 days. Across three separate experiments, a stool from the first week and a stool or cecal content sample from the third week was used to quantify *C. difficile* CFU per miligram (CFU/mg), via serial dilution onto *C. difficile* selective plates (Biomerieux), and TcdB levels per milligram of stool (TcdB/mg) using an ELISA kit (TCGbiomics). For the TcdB ELISAs, fecal samples in the microbiome engraftment screen were diluted 1:5 in dilution buffer, and manufacturer instructions were followed. To calculate ratios of CFU/mg or TcdB/mg for each mouse, the average level was determined across mice within each donor arm from the same experiment at an early timepoint (day 4 or 5) and the day 21 value was divided by this value. This was because some mice early after engraftment did not produce stool.

For the phylogenetic analyses of donor stool *C. difficile* isolates, a core-gene alignment between all 15 isolate genomes was generated using panaroo as above, and a tree was constructed using RAxML^55^. Average nucleotide identity (ANI) between closely related clade 1 isolate genomes was determined by running snp-dist(https://github.com/tseemann/snp-dists) with the panaroo core-gene alignment as input. ANI=(total number of basepairs in alignment-total number of SNPs)/ total number of basepairs in alignment.

### Statistical analyses of microbiome data

Species alpha diversity (richness and Shannon diversity) and the ARG richness and burden (RPKM) were compared across human cohort groups using the Wilcoxon rank sum test with Bonferroni correction for multiple hypotheses. Species beta diversity (Bray-Curtis distance matrix computed using vegdist) was compared between groups by PERMANOVA using adonis2 with 9999 permutations. In the human microbiome data, MetaPhlAn4 features were used in MaAsLin2^52^ to investigate differences between disease status or cephalosporin exposure using the formulas: Taxa∼ Group+(1 | Participant_ID) for all colonized patients or Taxa∼ ceph_30d+(1 | Participant_ID) for all hospitalized patients, respectively, with normalization=NONE and all other parameters set to default. Similarly, MetaPhlAn4 features (from all stool/cecal microbiomes from surviving mice, as filtered above) and HUMAnN3 features were used to understand microbiome correlates of *C. difficile* growth (CFU/mg) and toxin (TcdB/mg). The formula for the analyses in the microbiome engraftment screen was: Taxa/Pathways ∼ Growth + Timepoint +Toxin+(1 | DonorID) + (1 | Cdiffstrain) + (1 | CageAZ) + (1 | Sex) + (1 | MouseID), with min_prevalence=0.2, normalization=NONE, transform=LOG, and all other options used in default settings. The formula for the analyese in the defined community experiments to investigate community differences in metabolic pathway composition at day 28 was: Aggregated Pathways ∼ Community+ (1 | MouseID) + (1 | Cage) + (1 | Sex) + (1 | Strain). The q-value and coefficient were depicted in plots. Sparse estimates of correlations among microbiomes (SECOM) was used to investigate correlational relationships between species features^56^. Linear correlations of Metaphlan4 features was performed using secom_linear() with specifications: assay_name = "counts", tax_level = "Species", pseudo = 0,lib_cut=0, method = "pearson", soft = FALSE, thresh_len = 100, n_cv = 10, thresh_hard = 0, max_p = 0.005. P-value filtered correlations of interest were visualized using Cytoscape^57^.

### Defined community culturing

Stool gavage material used for animal experiments (from donors 4, 13, 15, and 16) was used to culture taxa representative of microbiomes of this patient population. For each donor, ethanol-treated stool^58^ was plated on LYBHI^59^ containing 0.1% sodium taurocholate, and non-ethanol-treated stool was plated on LYBHI in 10-fold dilutions. For ethanol-treated stool, stool was resuspended in an equal volume of 100% ethanol and left to sit for 4 hours. Following this incubation, serial dilutions were plated onto agar plates. Plates were incubated for 2-3 days; distinct colonies were subcultured into LYBHI in deep 96 well plates and allowed to grow for two days. Well contents were identified using V3-V4 16S rRNA sequencing^60^. Candidate taxa were subcultured to a clonal morphology and confirmed by 16S full-length Sanger sequencing via Genewiz. Purified clones were stored in LYBHI+25% glycerol at -80 degrees C.

### Defined community mouse experiments

All isolates were inoculated onto LYBHI agar plates for 3 days to allow growth. A subset of isolates were inoculated onto LYBHI+0.5% gastric mucin agar and grown in LYBHI+0.5% gastric mucin broth: both *Akkermansia muciniphila* strains, *Blautia hansenii, Alistipes putredinis, Anaerofustis stericorihominis*, *Anaerotruncus colihominis.* A single colony was inoculated into a deep-well plate with 1mL of LYBHI and diluted back 1:10 into fresh media every 24 hours for two days. Isolates were pooled at equivalent loads based on OD_600_, to achieve ∼10^9 CFU/mL in PBS. Germ-free adult (10-12 weeks of age) male and female C57BL-6J mice were gavaged once daily for two days with each community. Mice were followed over time for health (weight and clinical signs of distress). At 28 days, mice were gavaged with 10^4 CFU/mL of each *C. difficile* strain. Stool was also collected at Day 28 before infection and enumerated for total CFU/mg on LYBHI with 0.5% sodium taurocholate after 2 days of growth. Mice were assessed for health daily following gavage for seven days and euthanized as above. At days 2, 4, and 6, stool was collected and analyzed for *C. difficile* CFU/mg. Cecal content samples were collected following euthanasia. Toxin concentration (TcdB) in cecal samples was detected using ELISA kit for ‘Separate detection of *Clostridium difficile* TcdA or TcdB’ (tgcBIOMICS, Germany). Cecal samples were diluted 1:10 in dilution buffer and loaded onto ELISA plates for detection of toxin, as per the manufacturer’s instructions. TcdB levels were normalized to total protein in the corresponding cecal sample estimated by Bradford assay and expressed as ng/mg protein. For calprotectin estimation, cecal samples were diluted 1:5 in reagent diluent and quantified by ELISA using the DuoSet Mouse S100A8/S100A9 Heterodimer kit (R&D Systems, Minneapolis, MN, USA) according to the manufacturer’s recommendations. Calprotectin levels were normalized to total protein and expressed as pg/mg protein.

### Data analyses and visualization

All data distributions were depicted using RStudio or Prism and indicate the median with a line (whether in a box-and-whiskers plot or alone). Boxplots display the 25^th^ and 75^th^ percentile in the boxed region and depict the maximum and minimum value with the whiskers. Ellipses displayed on PCoA plots depict the 95% confidence interval for the specified group. All statistical tests were two-sided unless otherwise mentioned in the figure legends. All distributions were assessed for normality using the Shapiro-Wilkes test and then correspondingly tested for statistically significant differences using an experiment-specific parametric or non-parametric test.

## Data availability

Metagenomic data and its associated metadata is available at BioProject PRJNA1067813.

## Code availability

The random forest classifier code and accompanying antibiotic exposure information and metadata can be found at https://github.com/dantaslab/2024_Fishbein_DeVeaux_Cdiff_DefCom_Microbiome.

## Acknowledgements

The authors are grateful for members of the Dantas lab for their helpful feedback on the data analysis and preparation of the manuscript, with special thanks to Drew Schwartz and Mark Gorelik for their conceptual insights and Philip Frasse for assisting in sample preparation. The authors would also like to thank the Edison Family Center for Genome Sciences and Systems Biology staff, Eric Martin, Brian Koebbe, MariaLynn Crosby, and Jessica Hoisington-López for their expertise and support in sequencing/data analysis, Jacky Theodore, Bonnie Dee, James Anderson, and Keith Page for administrative support, and Sondra Seiler for her contributions to the prospective clinical study. Lastly, the authors express their gratitude towards Summer Rucknagel, J. Michael White, and Sian Madden from the Washington University School of Medicine Gnotobiotic Research, Education, and Transgenic Center for their valuable support in conducting the gnotobiotic mouse experiments described in this study. G.D. and ERD were supported in part by the National Institute of Allergy and Infectious Diseases (NIAID: https://www.niaid.nih.gov/) of the National Institutes of Health (NIH) under award number R01AI155893 (PI: G. Dantas) and an award from the Foundation for Barnes-Jewish Hospital and Institute of Clinical and Translational Sciences. This publication was also supported by the NIH/National Center for Advancing Translational Sciences (NCATS), grant UL1 TR002345 (PI: B. Evanoff). This work was also supported by funding through the CDC BAA #200-2018-02926 (PI: E. Dubberke). SRSF is supported by the National Institute of Child Health and Human Development (NICHD: https://www.nicdhd.nih/gov) of the NIH under award number T32 HD004010 (PI: P. Tarr) and through the NIAID of the NIH under award number K99AI17674 (PI: S. Fishbein). ALD is supported by the NIH under award number T32GM139774 (PI: H. True-Krob). The conclusions from this study represent those of the authors and do not represent positions of the funding agencies.

## Author contributions

SRSF, ALD, ALF, ERD, and GD conceived the study. SRSF, ALD, SK, ALF, JL, JN, MJW, and BM performed the gnotobiotic mouse work. SRSF, ALD, SK, JL, BM, and SL performed *in vitro* experiments and prepared sequencing data. SRSF, ALD, JL, WA, and SL prepared the defined communities with expert guidance from JG. SRSF, ALD, and WA performed computational analyses. SRSF and ALD wrote the manuscript with feedback from all authors. TH and KAR retrieved patient-associated metadata, provided patient stool samples, and cultured *C. difficile* isolates from primary human stools. All authors participated in reviewing and editing the manuscript. ERD and GD supervised the research.

---

**Supplementary Figure 1.**
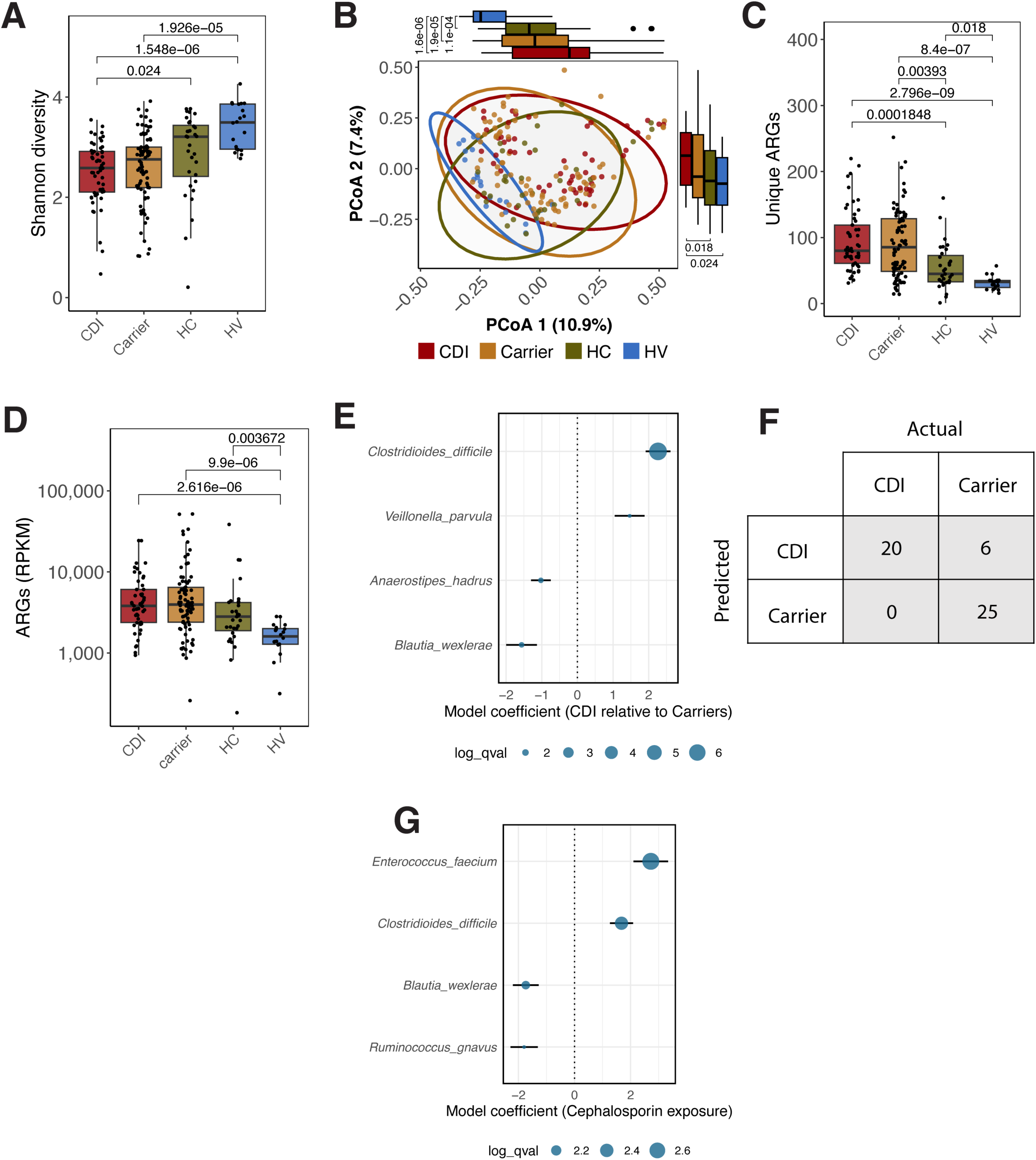
Microbiome features of *Clostridioides difficile* colonization at multiple scales. **A**) Taxonomic Shannon diversity index of stool metagenomes based on relative abundance of species using MetaPhlAn4. B) Principal coordinate analysis (PCoA) of Bray-Curtis distances inferred at the species level. Ellipses represent 95% confidence intervals for each clinical group. Percent variance explained by each axis shown is displayed parenthetically. C) Antimicrobial resistance gene (ARG) richness and D) burden (RPKM) within stool metagenomes based on shortBRED profiling. Statistical tests for A, C, D, and the marginal comparisons by axis in B were performed using the Wilcoxon rank-sum test with Bonferroni correction for multiple comparisons. E) Enrichment of species in patients with CDI relative to carriers using linear mixed effect models (MaAsLin2, as described in the Methods). Dot sizes correspond to Benjamini-Hochberg-corrected *p-*values (all false-discovery-rate [FDR] < 0.05) and standard deviations represented by bars. F) Confusion matrix for random forest classifier performance of the most accurate classifier (of 100 iterations). G) Species associations with recent cephalosporin exposure in colonized patients as carried out in E. HC, hospitalized control; HV, healthy volunteer. For A-D, N = 53 CDI, 82 carrier, 32 HC, 20 HV.

**Supplementary Table 1.** Antibiotic exposures in the previous 30 days for hospitalized patient groups. *P*-values determined across all three groups with Fisher’s exact test (*p* ≤ 0.05 bolded). HC, hospital control.

| Antibiotic exposures in the previous 30 days by class |  |  |  |  |
| --- | --- | --- | --- | --- |
|  | HC | Carrier | CDI | *p*-value |
| *n* | 32 | 82 | 53 |  |
| Any antibiotics (%) | 16 (50.0) | 18 (22.0) | 21 (39.6) | **0.006** |
| Intravenous vancomycin (%) | 8 (25.0) | 10 (12.2) | 21 (39.6) | **0.001** |
| Cephalosporin (%) | 3 (9.4) | 6 (7.3) | 29 (54.7) | **<0.001** |
| Carbapenem (%) | 4 (12.5) | 6 (7.3) | 8 (15.1) | 0.363 |
| Fluoroquinolone (%) | 6 (18.8) | 5 (6.1) | 11 (20.8) | **0.023** |
| Specific categories refer to classes with ≥10% prevalence. *p*-values determined by Fisher's exact test. |  |  |  |  |

**Supplementary Figure 2.**
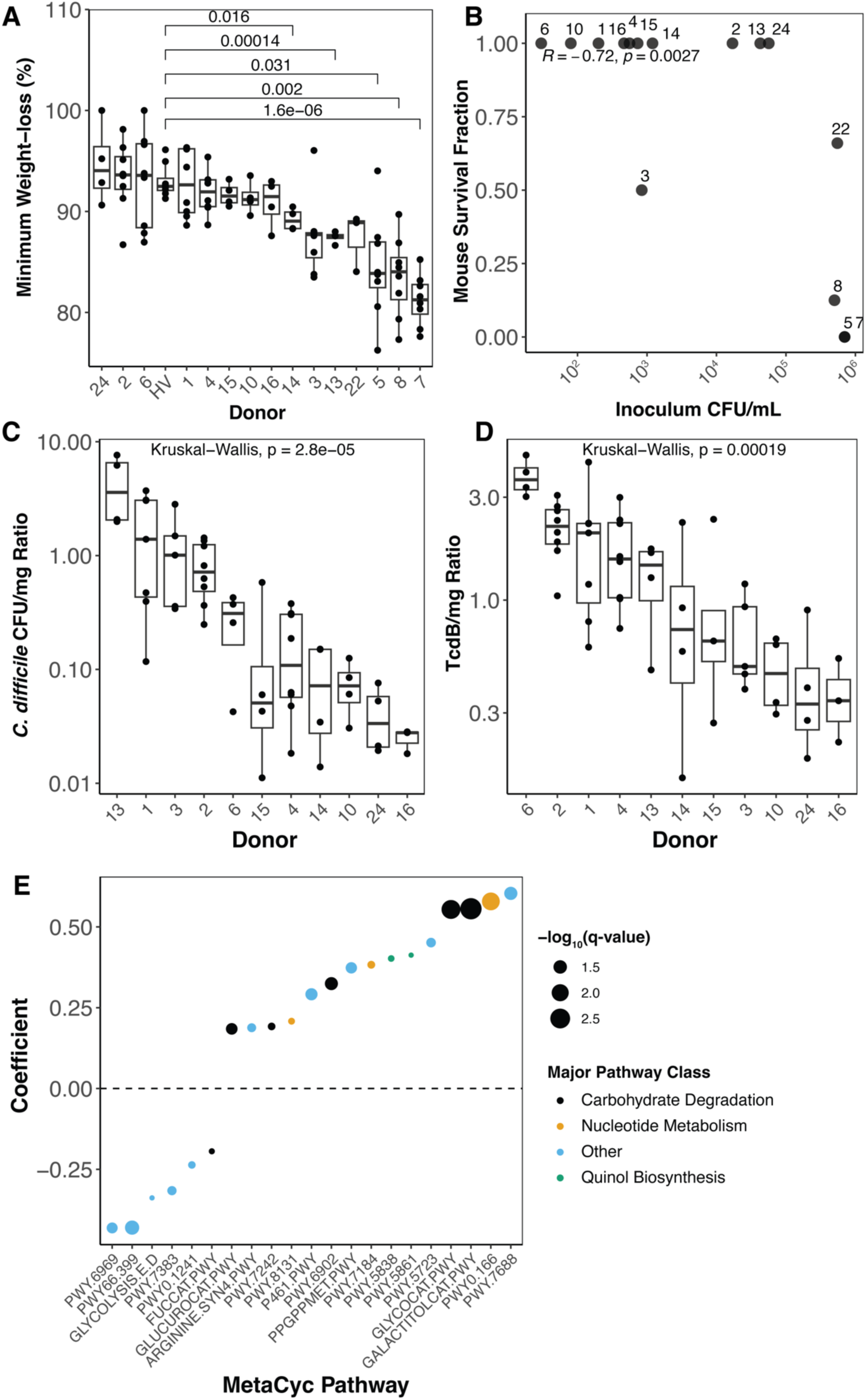
Donor-specific features of microbiome engraftment. A) Minimum weight loss across 21 days for each stool microbiome. Significant statistical differences for each donor relative to the control microbiome (Donor HV) as measured by multiple t-tests against Donor HV as a reference, with p-value adjustment performed by Bonferroni correction. B) Relationship between percentage of mice surviving over 21 days and *C. difficile* concentration in the inoculum. Boxplots of growth (C) and toxin (D) ratio of day 21 *C. difficile* levels relative to day 4-5 levels for each stool microbiome engrafted. E) Linear coefficients of MetaCyc pathways (HUMAnN3-produced) associated with increased *C. difficile* growth, q-value <0.1, as described in the Methods via MaAsLin2 (analyzed 88 stool metagenomes over time across 13 donor backgrounds)

**Supplementary Figure 3.**
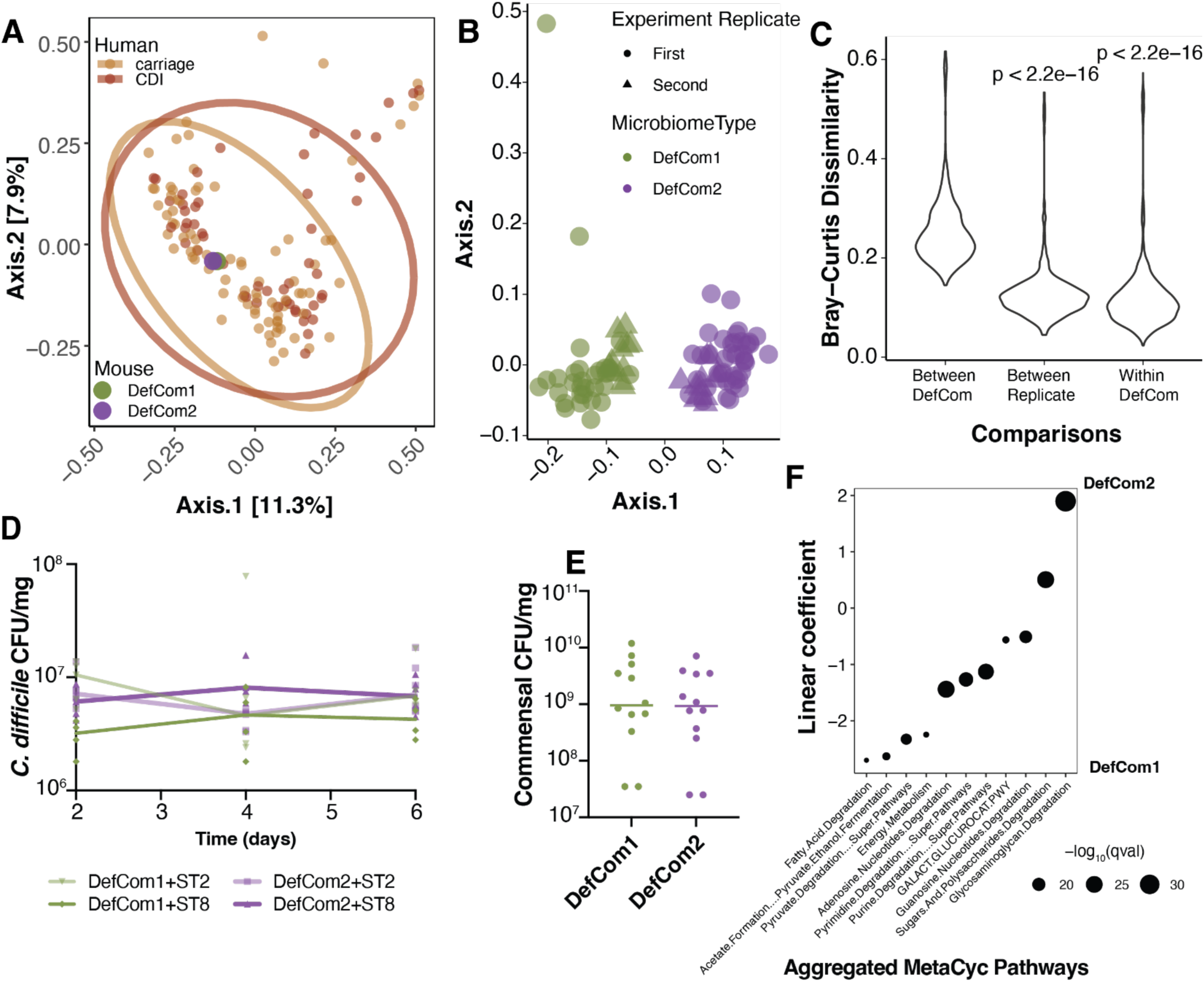
DefCom2 supports less virulent *C. difficile* proliferation. A) Principal coordinate analysis of Bray-Curtis dissimilarity between taxonomic features (aggregated to genus level) from human samples (small points) with representative DefCom1 and DefCom2 microbiomes (large points, averaged across all mice at day 28 [N=43 mice from DefCom1, N=52 mice from DefCom2]). B) Principal coordinate analysis of Bray-Curtis dissimilarity between mouse stool microbiomes from (A) at day 28 during each defined community experiment. C) Quantification of beta diversity (Bray Curtis dissimilarity) differences between different communities (Between DefCom), between experimental replicates within the same DefCom (Between Replicate), and between microbiomes from the same defined community across experimental replicates (Within DefCom). P-values indicated a Wilcoxon rank sum test performed comparing Between DefCom group to other groups. D) Stool levels of *C. difficile* post-infection (left, from Figure 4E) with area under the curve (AUC, right panel), measured across all three time points for ST8 infection in DefCom1 (N=5 mice) and DefCom2 (N=6). **, p<0.01, as obtained by Mann-Whitney test. E) Stool levels of commensal bacteria in each defined community at day 28 of second experiment (N= before *C. difficile* infections. F) Linear coefficients of MetaCyc-aggregated pathway differences between DefCom1 and DefCom2 microbiomes, q-value<1e-15 are depicted.

**Supplementary Table 2.**
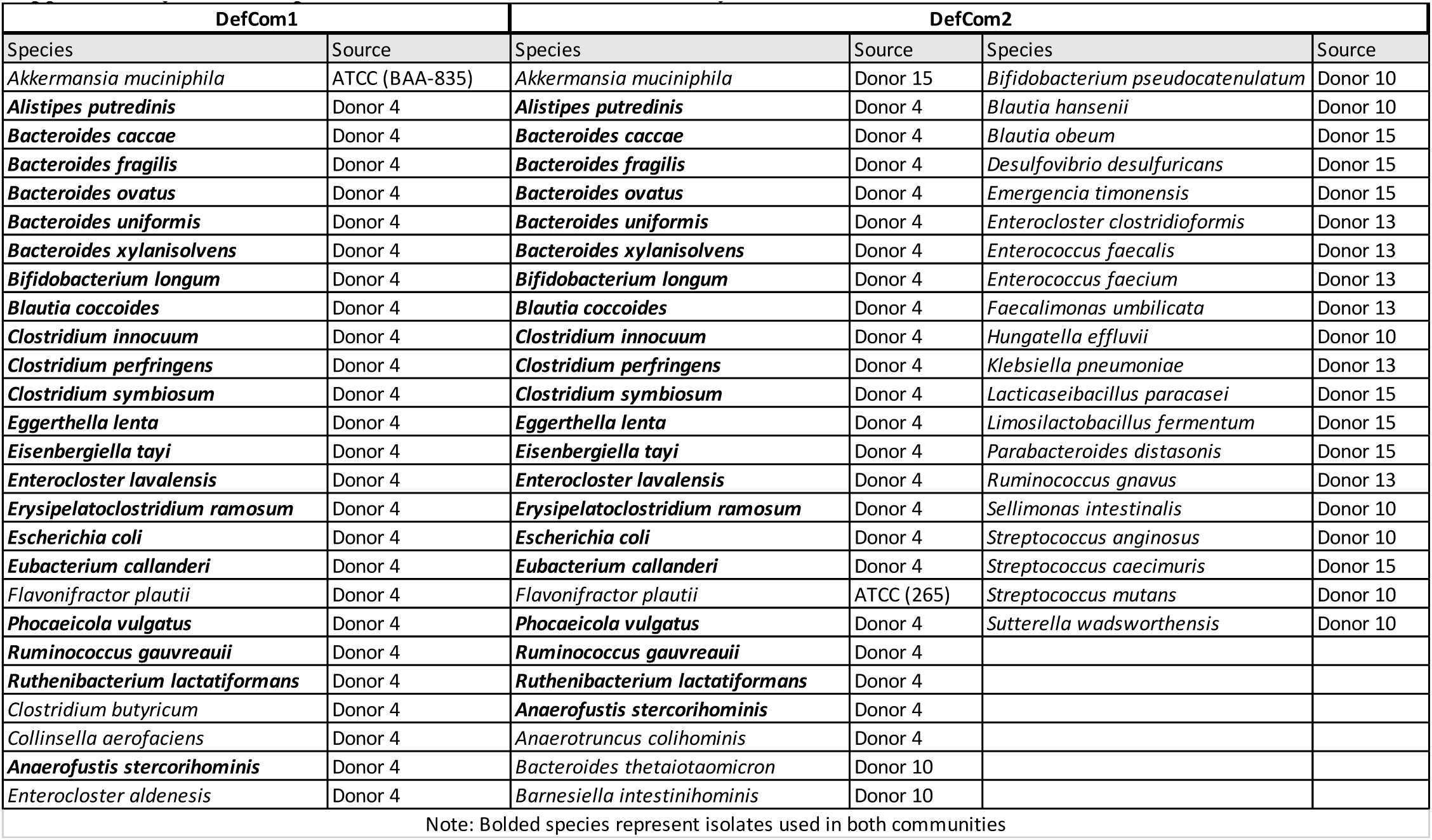
Species used in each defined community.

